# Tissue-Restricted Inhibition of mTOR Using Chemical Genetics

**DOI:** 10.1101/2022.03.08.483534

**Authors:** Douglas R. Wassarman, Kondalarao Bankapalli, Leo J. Pallanck, Kevan M. Shokat

**Affiliations:** Department of Cellular and Molecular Pharmacology, Howard Hughes Medical Institute, University of California San Francisco, San Francisco, California 94158, United States; Department of Genome Sciences, University of Washington, Seattle, Washington 98195, United States

**Keywords:** mTOR, rapamycin, kinase inhibitor, tissue specific, Drosophila

## Abstract

mTOR is a highly conserved eukaryotic protein kinase that coordinates cell growth and metabolism and plays a critical role in cancer, immunity, and aging. It remains unclear how mTOR signaling in individual tissues contributes to whole-organism processes because mTOR inhibitors, like the natural product Rapamycin, are administered systemically and target multiple tissues simultaneously. We developed a chemical-genetic system, termed selecTOR, that restricts the activity of a Rapamycin analog to specific cell populations through targeted expression of a mutant FKBP12 protein. This analog has reduced affinity for its obligate binding partner FKBP12, which reduces its ability to inhibit mTOR in wild-type cells and tissues. Expression of the mutant FKBP12, which contains an expanded binding pocket, rescues the activity of this Rapamycin analog. Using this system, we show that selective mTOR inhibition can be achieved in *S. cerevisiae* and human cells, and we validate the utility of our system in an intact metazoan model organism by identifying the tissues responsible for a Rapamycin-induced developmental delay in *Drosophila*.

**Significance Statement:** mTOR plays a number of critical organismal roles, including in cell growth, development, immunity and aging, but dissecting the tissue-specific influences of mTOR has proven challenging. This work describes a simple system for identifying the specific tissues and cells responsible for the diverse functions of mTOR, and we show that our system can be used in organisms ranging from yeast to humans.

## Introduction

Systemic inhibition of the conserved energy sensing kinase mTOR by the natural product Rapamycin leads to a number of physiological effects with significant therapeutic potential (1, 2). Rapamycin is used clinically as an immunosuppressant to prevent organ transplant rejection and in the treatment of breast and kidney cancer, and it is under investigation to combat aging (3, 4). However, the beneficial effects of Rapamycin remain coupled to toxicities that limit its broader clinical application including insulin resistance (5, 6) and impaired wound healing (7), which result from on-target inhibition of mTOR.

Although these beneficial and adverse effects result from inhibition of the same molecular target, many do not derive from the same cell types, making it possible to isolate them through tissue-specific inhibition of mTOR. Genetic disruption of the mTOR pathway in skeletal muscle causes insulin resistance in mice (8), but disruption in adipose increases insulin sensitivity (9). Inhibition of mTOR prevents differentiation of T helper cell lineages but stimulates generation of memory T cells (10, 11). In *Drosophila*, genetic inhibition of TOR signaling in adipose is sufficient to extend lifespan (12). The outcomes of systemic mTOR inhibition are the union of these varying, and sometime opposing, effects in different cell types and tissues.

In order to understand the organismal effects of Rapamycin, it is necessary to investigate the tissue-specific effects of pharmacological mTOR inhibition, rather than rely solely on genetic models, as the outcomes of pharmacological inhibition often differ from those of genetic knockout (13). This is especially true with Rapamycin, which acts as a molecular glue between the peptidyl prolyl isomerase FKBP12 and the mTOR FRB domain to inhibit phosphorylation of only a subset of mTOR complex 1 (mTORC1) substrates (14), and inhibit or activate mTOR complex 2 (mTORC2) depending on cell type and treatment duration (15). This mode of inhibition is difficult to replicate genetically and yet is the most critical to understand for therapeutic applications.

Here we present selecTOR, a chemical-genetic platform that restricts the activity of a Rapamycin analog to specific cell populations through targeted expression of a mutant FKBP12 protein. This analog has decreased affinity for FKBP12, rendering it ineffective at binding and inhibiting mTOR, but preferentially binds a mutant FKBP12 with an expanded pocket, allowing it to selectively inhibit mTOR in cells where this mutant protein is expressed. We show that selecTOR can be used to restrict mTOR inhibition in multiple cell types, and we validate the utility of this system in *Drosophila* by investigating the roles of the musculature, the nervous system, and the ring gland in Rapamycin-mediated developmental delay.

## Results

Rapamycin’s unusual mechanism of action presented an opportunity to combine chemical and genetic modifications to target mTOR in specific cell populations. Rapamycin alone has low affinity for mTOR (16). It is the Rapamycin-FKBP12 complex that presents a complementary binding surface for the mTOR FRB domain to assemble into a ternary complex. Therefore, we predicted that modifying Rapamycin such that it could no longer bind FKBP12 would render it ineffective at inhibiting mTOR and that restoring FKBP12-binding of this analog through mutation of the FKBP12 binding site would, in turn, rescue formation of the inhibitory ternary complex (Figure 1a). Even when administered systemically, this Rapamycin analog would inhibit mTOR only in cell populations engineered to express the mutant FKBP12.

**Figure 1.**
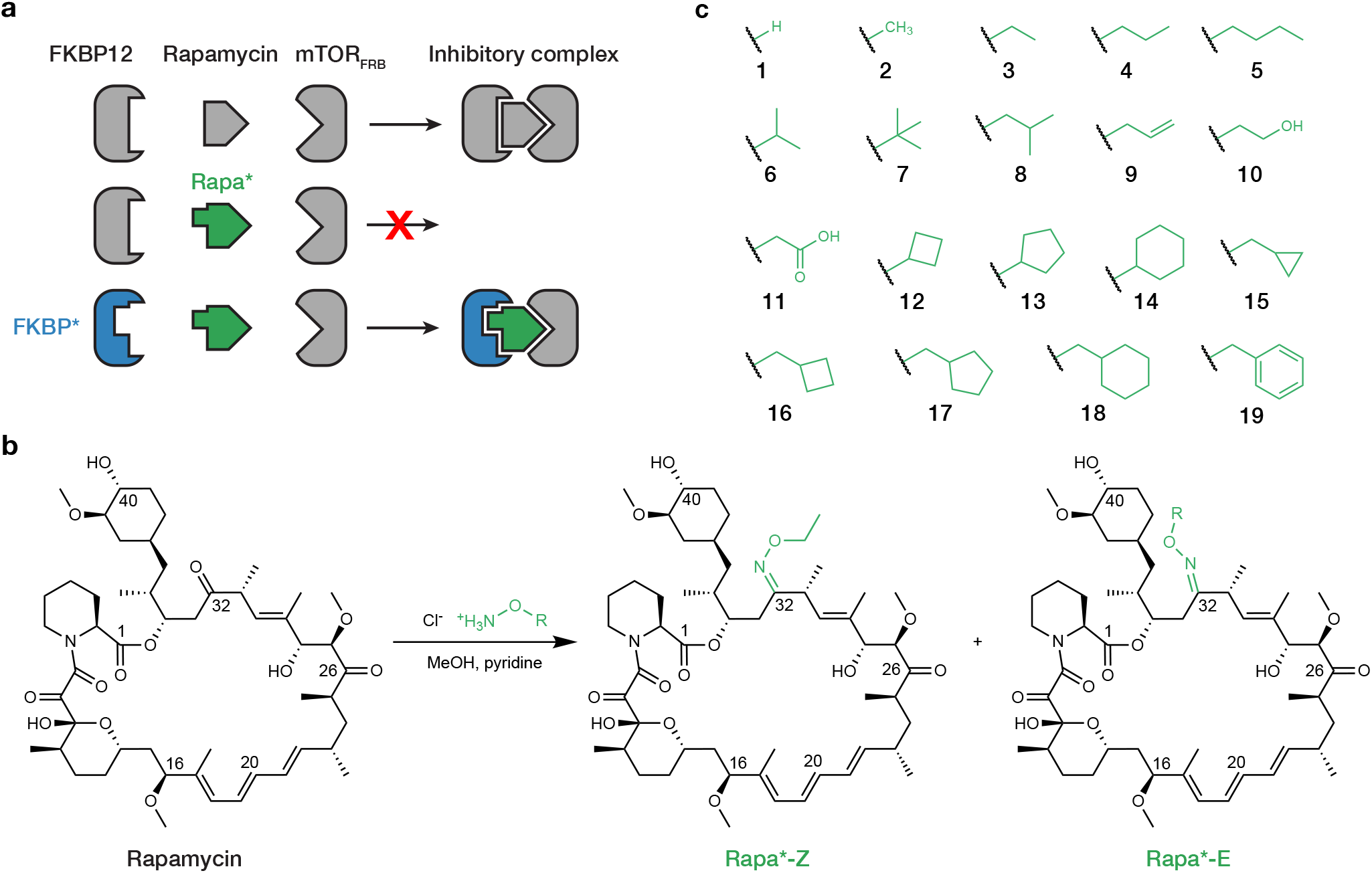
Design and synthesis of an orthogonal Rapamycin analog. (a) Rapamycin forms an inhibitory complex by binding FKBP12 and mTOR FRB domain. Rapa*, a “bumped” analog of Rapamycin, does not bind wild-type FKBP12 but does bind the “hole” mutant FKBP*. (b) Synthetic scheme for oxime ligation of Rapamycin to generate Rapa* *Z-* and *E*-isomers. (c) Assembled Rapa* library containing various oxime R-groups.

We first set out to design Rapamycin analogs with reduced affinity for FKBP12. Reactions have been reported for direct modification of Rapamycin at C-16, C-20, C-32, and C-40 (17–20). Crystal structures of the Rapamycin-FKBP12 complex show that the C-32 ketone is the only position located inside the binding pocket, where it is closely flanked by the side chains of F46 and V55 (21). Reaction of Rapamycin with an *O*-functionalized hydroxylamine generated a separable mixture of the corresponding *E-* and *Z*-oxime at the C-32 position (Figure 1b). We constructed a library of 38 (19 substituents x 2 isomers) Rapamycin analogs, or Rapa* compounds, with varying size, geometry, and hydrophobicity to probe the stringency of the FKBP12 binding site (Figure 1c).

We selected *S. cerevisiae* as a model to test the Rapamycin analogs because of its exquisite growth sensitivity to Rapamycin and the high conservation of its mTOR and FKBP12 orthologs. Knockout of *fpr1*, the yeast ortholog of FKBP12, conferred resistance to Rapamycin in a disk diffusion assay, and introduction of human FKBP12 rescued full sensitivity (22) (Figure 2a). We also introduced a series of FKBP12 mutants with substitutions to smaller amino acids at F46 or V55 to create space in the binding pocket to accommodate the additional bulk of the Rapamycin analogs. We expected that these mutations would be tolerated because it has been shown previously that mutations to F46 or V55 have only a modest effect on FKBP12 rotamase activity (23). Immunoblots against a C-terminal HA-tag showed that expression of FKBP12 mutant proteins was reduced compared to wild-type FKBP12 (Figure 2b). This suggests that the mutant proteins are somewhat destabilized despite their reported enzymatic activity.

**Figure 2.**
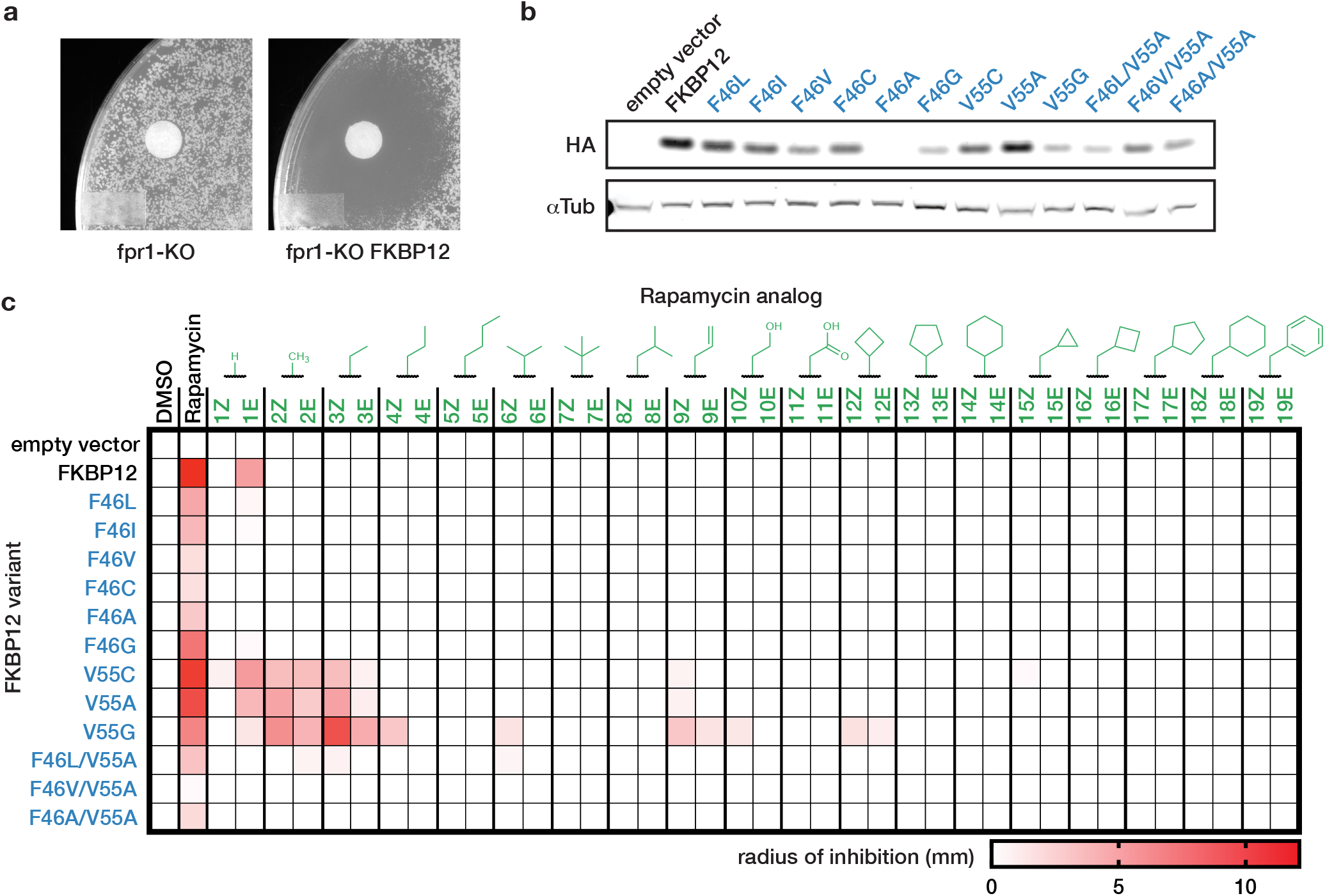
Mutant FKBP12 expression sensitizes *S. cerevisiae* to inhibition by Rapa*. (a) Susceptibility of *S. cerevisiae* BY4741 fpr1-KO with empty vector or vector encoding human FKBP12 to 100 μM Rapamycin after 3 days. (b) Expression of C-terminal HA-tagged human FKBP12 variants in fpr1-KO yeast. (c) Growth inhibition of fpr1-KO yeast expressing FKBP12 variants by 100 μM of Rapa* compounds. Radius of inhibitory zone shown in mm.

We determined the sensitivity of the different FKBP12-expressing yeast strains to the panel of Rapamycin analogs by measuring the zone of growth inhibition caused by each compound in a disk diffusion assay (Figure 2c). Nearly all Rapamycin analogs failed to inhibit growth in the wild-type FKBP12 strain, indicating that even small oximes at C-32 are sufficient to disrupt FKBP12 binding and prevent mTOR inhibition. A subset of FKBP12 mutants rescued sensitivity to some of the Rapamycin analogs. The combination of Rapa*-3Z and FKBP12 V55G displayed the highest level of inhibition, similar to that of the Rapamycin and wild-type FKBP12 combination. Cyclic and large, branched analogs were poorly tolerated across the panel of mutants, while short linear substituents exhibited activity exclusively with the V55 mutants. In each case, the *Z* isomer outperformed its *E* counterpart, suggesting that the geometry of this isomer is more closely complemented by the pocket created by mutating V55. Interestingly, Rapamycin retained activity, albeit reduced, in the mutant strains, which indicates that the mutations did not preclude Rapamycin binding or ternary complex formation.

We next sought to quantify and further characterize the selectivity of the Rapamycin analogs for mutant FKBP12 biochemically. Using Time Resolved Fluorescence Resonance Energy Transfer (TR-FRET), we measured the drug-induced association of mTOR FRB with wildtype or mutant FKBP12 (Figure 3a). Each of the Rapamycin analogs that displayed high FKBP12 V55G selectivity in yeast also selectively induced the mutant ternary complex *in vitro*, whereas Rapamycin was slightly selective towards the wild-type complex (Supplemental Figure S1). The control compound FK506, which binds FKBP12 but not mTOR, failed to induce either ternary complex (Supplemental Figure S1). Rapa*-3Z once again emerged with the most favorable combination of potency and mutant selectivity, inducing the FKBP12 V55G ternary complex with an EC_50_ of 11.8 nM and the wild-type complex with an EC_50_ of 639 nM (Figure 3b).

**Figure 3.**
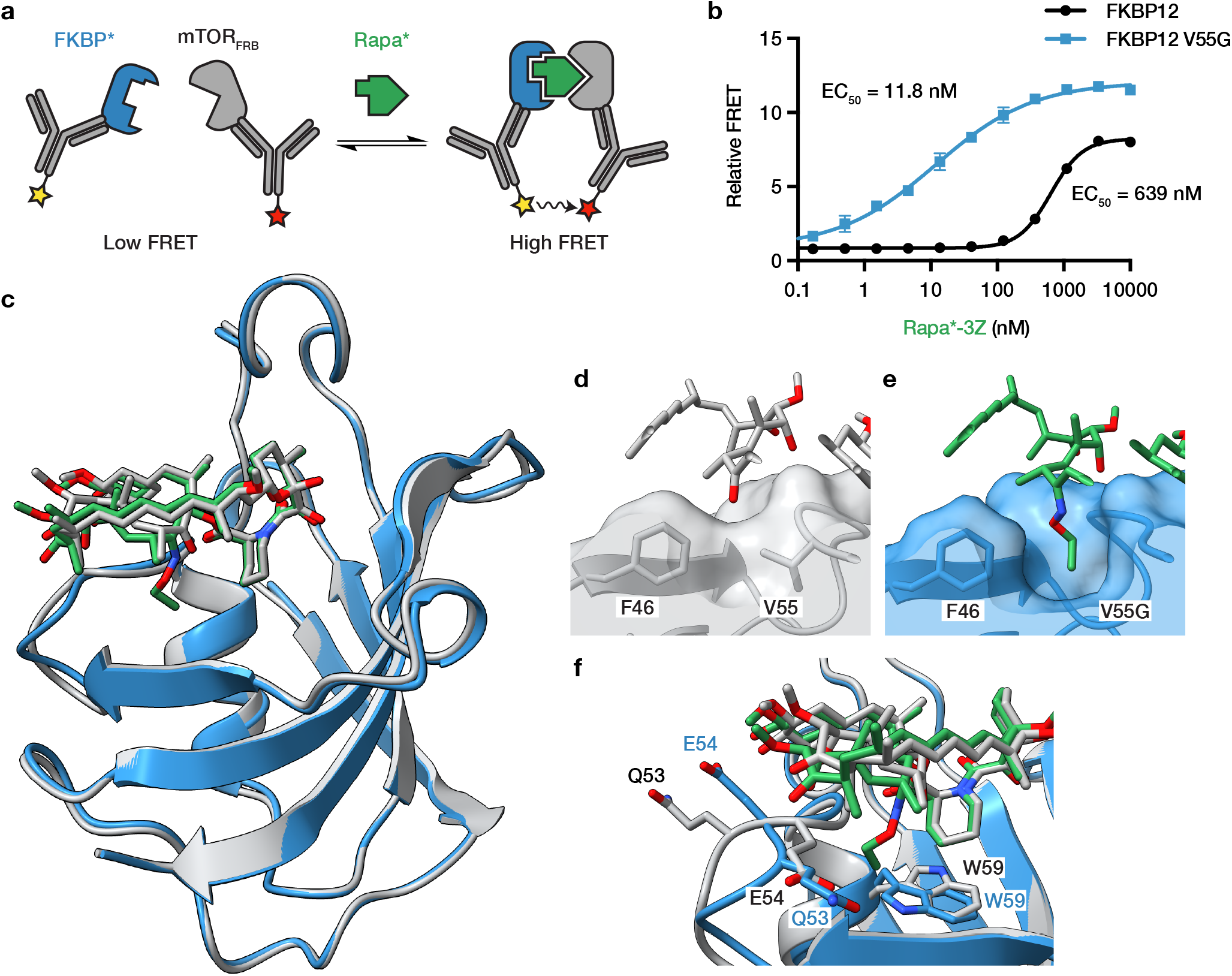
Rapa*-3Z selectively binds FKBP12 V55G. (a) Time-Resolved Fluorescence Resonance Energy Transfer (TR-FRET) reports association of recombinant HA-tagged wild-type or V55G mutant FKBP12 with GST-tagged mTOR FRB domain. (b) Ternary complex induction by Rapa*-3Z. n = 3, error bars show s.d. (c) X-ray crystal structure of Rapa*-3Z (green) bound to FKBP12 V55G (blue) aligned with a published structure of Rapamycin (grey) bound to FKBP12 (grey), PDB: 1FKB. (d) Rapamycin C-32 ketone and FKBP12 surface. (e) Rapa*-3Z C-32 ethyloxime and FKBP12 V55G surface. (f) Rearrangement of Q53, E54, and W59 residues between Rapamycin/FKBP12 and Rapa*-3Z/FKBP12 V55G structures.

To understand the mechanism of this selectivity at the atomic level, we solved a high-resolution X-ray crystal structure of Rapa*-3Z bound to mutant FKBP12 (Supplemental Table S1, PDB: 7U8D). The asymmetric unit contained two protein molecules which aligned closely with one another (RMSD = 0.3 Å) and are each bound to one Rapa*-3Z molecule (Supplemental Figure S2). The Rapa*-3Z molecules occupy similar overall conformations except for a 180° rotation of the cyclohexyl group, which suggests there is some flexibility in this region even when bound (Supplemental Figure S2). Our structure aligned closely with a published structure of Rapamycin bound to FKBP12 (21) (RMSD = 0.4 Å), an indication that the modifications did not significantly alter the overall protein fold or binding pose (Figure 3c). In the wild-type structure, the C-32 ketone of Rapamycin points toward the F46 and V55 side chains (Figure 3d). In our structure, the added ethyloxime moiety extends into a pocket created by the V55G mutation (Figure 3e). We see that this substituent clashes with the wild-type residue, preventing the binding of Rapa*-3Z and leading to its selectivity for the mutant protein.

The mutant structure also revealed conformational changes to nearby residues Q53, E54, and W59 (Figure 3f). Q53 and E54, which make up part of the loop that packs under Rapamycin C-28, have rotated and swapped positions in the mutant structure, leading to a closer interaction between the alcohol on Rapa*-3Z and the π-system of E54. It may be the additional flexibility from the nearby glycine mutation that allows this loop to flip and occupy this new arrangement. We also see that the indole side chain of W59 rotates in the mutant structure to meet the ethyl group of Rapa*-3Z, helping to fill the void left by the V55G mutation. Taken together, these observations show that it is both the creation of a larger binding pocket and subtle restructuring of the surrounding region that allow FKBP12 V55G to tightly bind Rapa*-3Z.

We next determined if Rapa*-3Z could be used to selectively inhibit mTOR in human cells expressing mutant FKBP12. Achieving selectivity in human cells is more challenging due to the fact that the human genome encodes 14 different FKBP proteins (24). Most of these family members are known to bind Rapamycin (25) and several may contribute to Rapamycin-dependent mTOR inhibition (26). Using the Flp-In T-REx system, we generated a 293 cell line with inducible over-expression of HA-tagged FKBP12 V55G. We treated this mutant cell line and the parental cell line with Rapa*-3Z for 24 hours and measured mTOR pathway outputs by Western blot (Figure 4a). Rapa*-3Z inhibited mTOR signaling selectively in the mutant cells over a wide dose range. Phospho-S6K and phospho-S6, both downstream of mTORC1, were inhibited with IC_50_’s of 1.0 nM and 1.3 nM, respectively, in the mutant cells compared to 76 nM and 121 nM in the wild-type cells (Figure 4a and Supplemental Figure S3). This shows nearly a 100-fold increase in sensitivity in the mutant-expressing cells. Phosphorylation of the mTORC2 substrate, Akt, was less sensitive overall to inhibition, a phenomenon that has also been observed with Rapamycin (27, 28), but Rapa*-3Z selectively reduced Akt phosphorylation in the mutant 293 cells at higher doses (Figure 4a).

**Figure 4.**
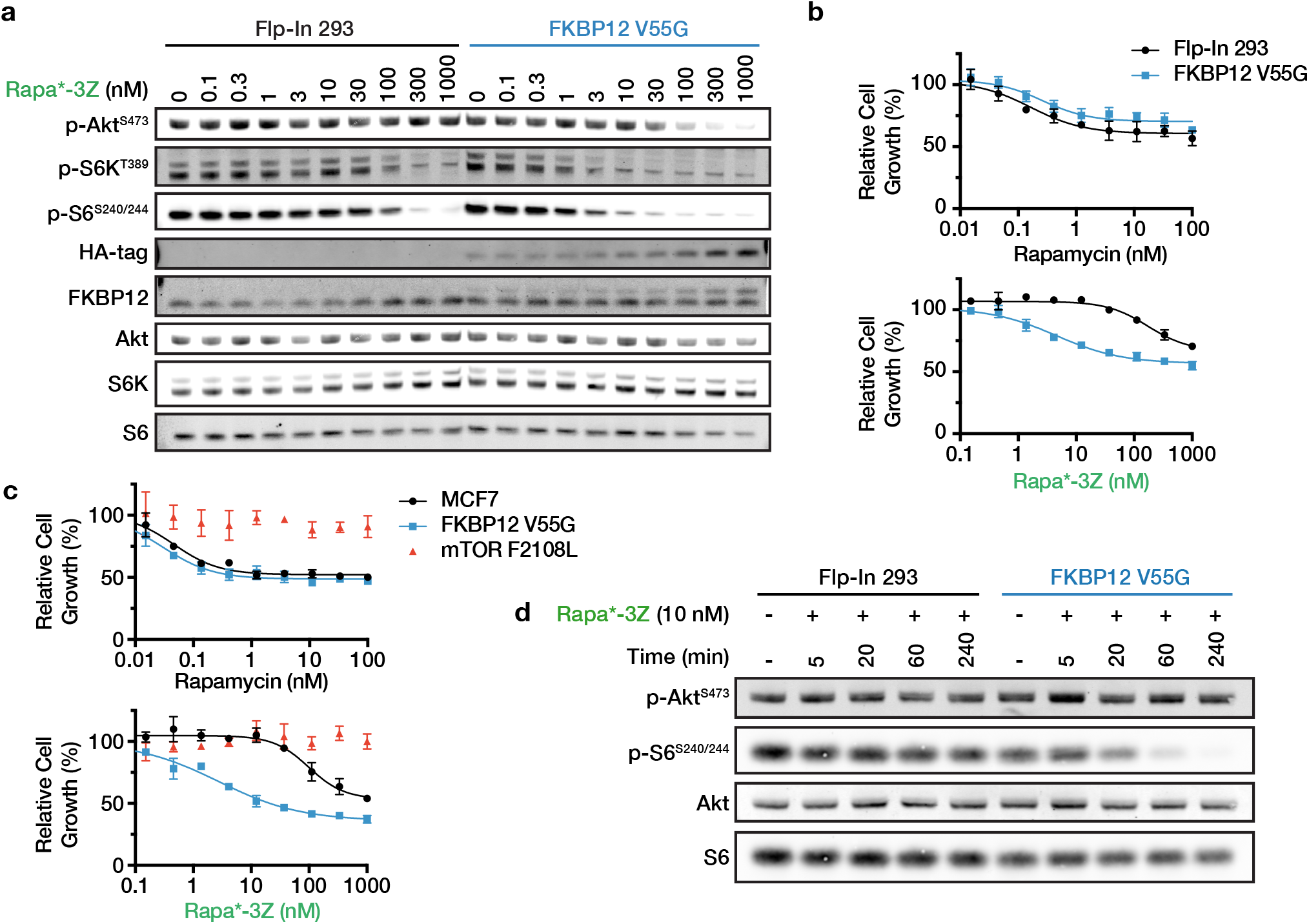
Rapa*-3Z selectively inhibits cultured human cells expressing FKBP12 V55G. (a) mTOR signaling following 24h treatment with Rapa*-3Z in Flp-In 293 cells with or without FKBP12 V55G-HA expression. (b) Cell growth determined by CellTiter-Glo after 72h Rapa*-3Z treatment in Flp-In 293 cells with and without FKBP12 V55G-HA expressing cells. (c) Cell growth determined by CellTiter-Glo after 72h Rapa*-3Z treatment in MCF7 cells with and without FKBP12 V55G-HA expression, and with the Rapamycin resistance mutation mTOR F2108L. Cell growth was normalized to DMSO control. n = 3, error bars show s.d. (d) Onset of mTOR inhibition following 10 nM Rapa*-3Z treatment in Flp-In 293 cells with and without expression of FKBP12 V55G.

Interestingly, we observed ligand-dependent stabilization of FKBP12 V55G (Figure 4a). A C-terminal HA-tag allowed us to detect expression of the mutant protein and also decreased the protein’s electrophoretic mobility enough to resolve it from wild-type FKBP12. Mutant expression was well below that of wild-type FKBP12 in the DMSO-treated cells, but in Rapa*-3Z treated cells expression increased dramatically. This also supports the conclusion that the V55G mutation is somewhat destabilizing to FKBP12. However, even low levels of FKBP12 V55G appear to be sufficient to sensitize cells to inhibition by Rapa*-3Z.

Due to mTOR’s central role in cell growth, its inhibition causes a decrease in cell proliferation. We observed that Rapa*-3Z selectively inhibited growth of mutant-expressing 293 cells after 72-hour treatment, while Rapamycin inhibited mutant and wild-type cells equally (Figure 4b). This result was replicated in the breast cancer cell line MCF7, where Rapa*-3Z activity was restricted to the mutant FKBP12-expressing cells (Figure 4c). To confirm that the observed growth inhibition was mediated through mTOR, we also tested Rapa*-3Z on MCF7 cells that harbored an mTOR mutation in the FRB domain, F2108L, that confers resistance to Rapamycin-based inhibitors (29, 30). Rapa*-3Z had no effect on the growth of these cells (Figure 4c).

One advantageous feature of small molecules is their ability to rapidly inhibit a protein target. This is especially important for signaling kinases, like mTOR, which are often regulated on the order of minutes rather than hours or days. After treatment with 10 nM Rapa*-3Z, mTOR signaling was rapidly inhibited in mutant FKBP12 293 cells. The t_1/2_ for inhibition of S6 phosphorylation was 31 minutes (Figure 4d and Supplemental Figure S4). At this dose, the wildtype 293 cells were unaffected, demonstrating that mutant FKBP12 expression enables specific and rapid inhibition of the mTOR pathway by Rapa*-3Z.

To evaluate the utility of this selecTOR system in an intact model organism, we placed the human FKBP12 V55G coding sequence downstream from a GAL4-responsive Upstream Activating Sequence (UAS) and used this construct to create transgenic flies (Figure 5a). We then examined the influence of Rapa*-3Z on a developmental defect caused by TOR pathway inhibition. Previous work has shown that Rapamycin treatment dramatically delays or abolishes larval development through inhibition of TOR pathway activity (31–33). As expected, we also observed profound inhibition of larval development upon Rapamycin treatment of larvae (Figure 5b and 5c). However, we detected no effect of Rapa*-3Z on larval development in wild-type animals, even at high concentrations (Supplemental Figure S5). Ubiquitous expression of the FKBP12 V55G protein also had no apparent effect on larval development in the absence of Rapa*-3Z (Figure 5b, 5c, and Supplemental Figure S6). However, Rapa*-3Z treatment of transgenic animals ubiquitously expressing the FKBP12 V55G protein potently inhibited larval development.

**Figure 5.**
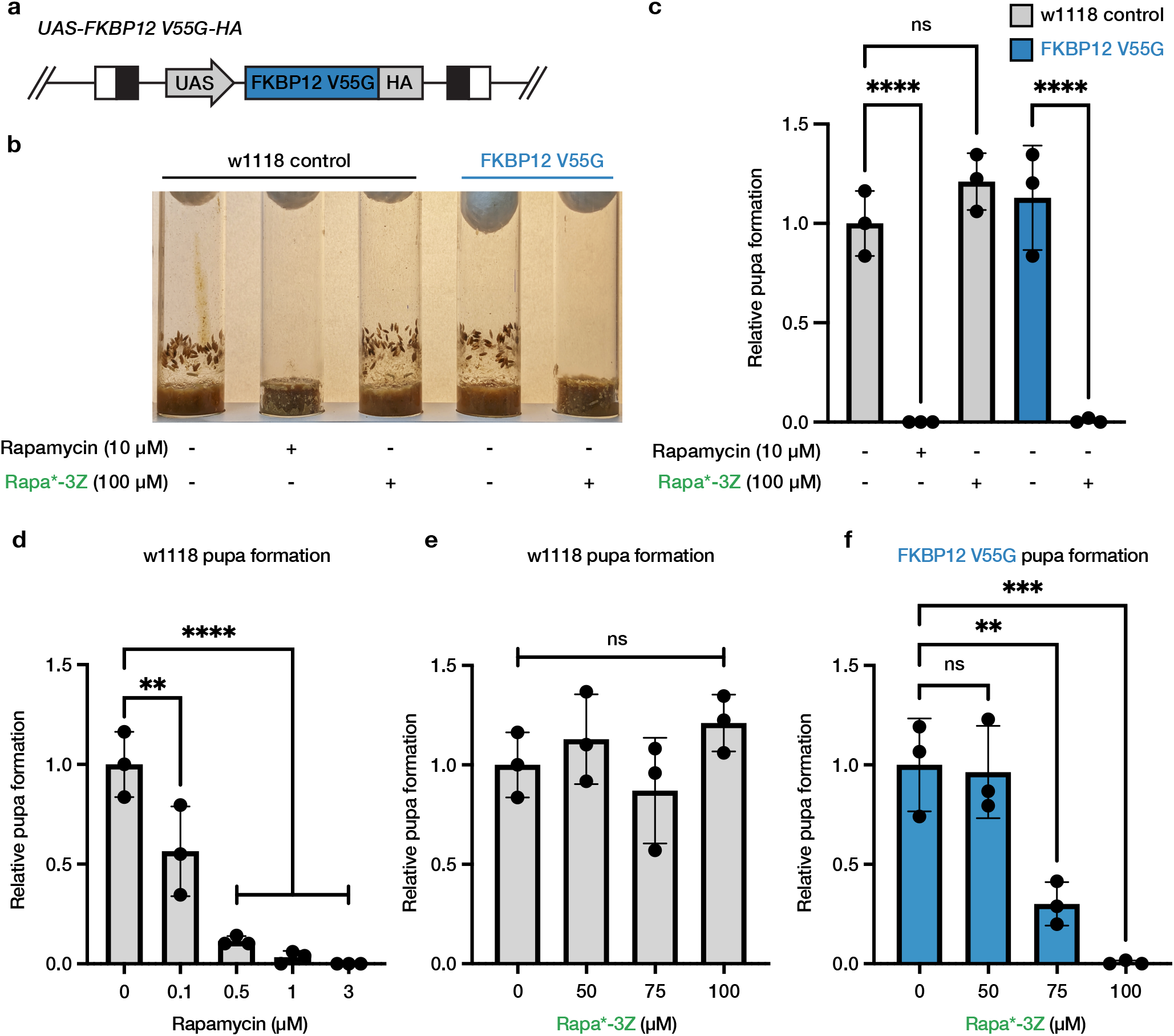
Rapa*-3Z inhibits larval development in transgenic animals expressing mutant FKBP12 V55G. (a) Schematic diagram of the GAL4-responsive FKBP12 V55G transgene. (b) Vials illustrating the number of control (w1118) and transgenic animals (FKBP12 V55G) that develop to the pupal stage in the presence and absence of 10 μM Rapamycin or 100 μM Rapa*-3Z. Images were captured 9 days after egg laying. (c) The relative number of pupae detected 9 days after egg laying in animals of the indicated genotypes in the presence and absence of 10 μM Rapamycin or 100 μM Rapa*-3Z. (d-f) Dose response effect of Rapamycin (d) and Rapa*-3Z (e and f) on the development of control (w1118; d and e) and transgenic (FKBP12V55G; f) larvae 9 days after egg laying. At least three independent experiments were performed (n=3). Significance was determined by one-way ANOVA multiple comparison test * p < 0.05; ** p < 0.001; *** p = 0.0001; **** p < 0.0001.

To directly compare the effective dose of Rapa*-3Z on larval development with that of Rapamycin, we performed a dose-response experiment (Figure 5d-5f). This work indicated that the dose of Rapa*-3Z necessary for complete inhibition of larval development in FKBP12 V55G animals was 100 μM, while 1 μM of Rapamycin was required to achieve complete inhibition in wild-type flies. This difference in potency is likely due to the decreased relative permeability and bioavailability of Rapa*-3Z as observed by parallel artificial membrane permeability assay (PAMPA) and Caco-2 permeability measurements (Supplemental Figure S7). Further experiments indicate that Rapa*-3Z is fully stable under the conditions of our experiments for at least 15 days (Supplemental Figure S8). Moreover, treating wild-type larvae with Rapa*-3Z after prolonged storage of the compound under experimental conditions had no apparent effect on larval development, indicating that Rapa*-3Z does not decompose to Rapamycin under these conditions (Supplemental Figure S8).

Previous work on the influence of Rapamycin on larval development suggested that the cause of the larval developmental delay could originate in the musculature, the nervous system, and/or the ring gland, where the larval molting hormone ecdysone is produced (31–33). To address this matter, we expressed FKBP12 V55G using *GAL4* drivers specific for one of these three tissues and then treated the animals with Rapa*-3Z. Specifically, we used the muscle-specific *mhc-GAL4* driver, the pan-neuronal *nSyb-GAL4* driver, and the ring-gland-specific *phm-GAL4* driver (Figure 6a). We treated these animals with 100 μM Rapa*-3Z, the minimal dose that completely prevented pupal formation 9 days after egg laying when FKBP12 V55G was expressed ubiquitously. We found that animals expressing FKBP12 V55G in the ring gland failed to pupate 9 days after egg laying in the presence of Rapa*-3Z (Figure 6b and 6c). We detected similar, but less severe, inhibition of pupal formation in Rapa*-3Z-treated animals expressing FKBP12 V55G in the musculature and nervous system. To verify these findings, we repeated this work using an independent collection of GAL4 drivers specific for the ring gland (*2-286-GAL4*), musculature (*Mef2-GAL4*), and nervous system (*elav-GAL4*). Consistent with our previous results, we found that expression of FKBP12 V55G in the ring gland nearly eliminated the larval to pupal transition in the presence of Rapa*-3Z, with significant, but lesser, inhibition conferred by expression of FKBP12 V55G in the musculature and nervous system (Supplemental Figure S9). Thus, our findings suggest that Rapamycin’s effects on *Drosophila* larval development are mediated primarily by TOR pathway inhibition in the ring gland, but also in the musculature and the nervous system.

**Figure 6.**
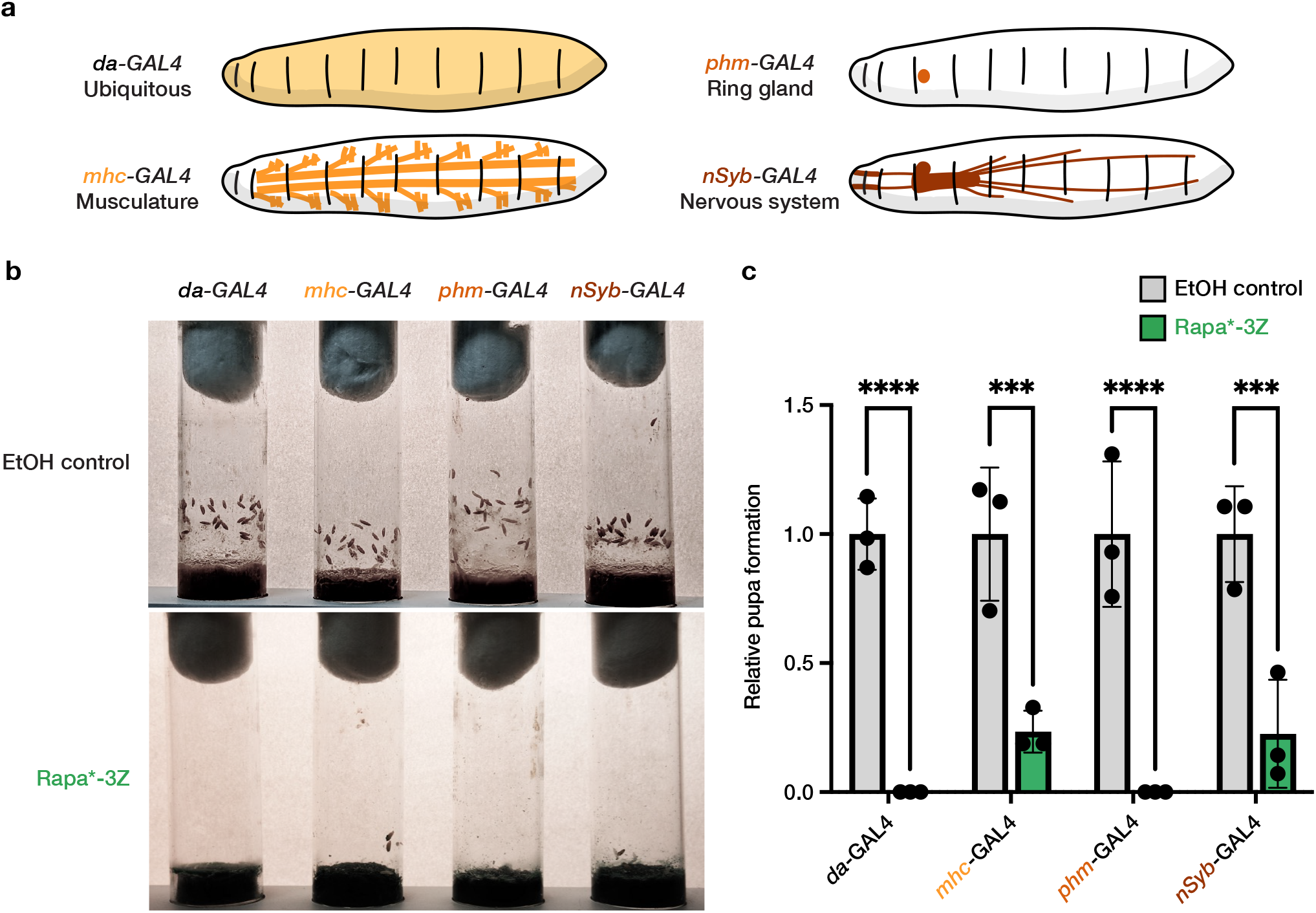
TOR pathway inhibition in ring gland, musculature, and nervous system delayed larval development in flies. (a) Schematic representation of FKBP12 V55G expression, ubiquitous (*da-GAL4*), musculature (*mhc-GAL4*), ring gland (*phm-GAL4*), or nervous system (*nSyb-GAL4*). (b) Vials illustrating the number of animals of the indicated genotypes that develop to the pupal stage in the presence or absence of 100 μM Rapa*-3Z 9 days after egg laying. (c) The relative number of pupae detected 9 days after egg laying in animals of the indicated genotypes treated with 100 μM Rapa*-3Z (n=3). Significance was determined by group comparison via two-way ANOVA * p < 0.05; ** p < 0.001; *** p = 0.0001; **** p < 0.0001.

## Discussion

Rapamycin remains a promising therapeutic across broad indications including cancer and aging, but in many instances, the tissues responsible for the beneficial effects of Rapamycin are unknown and unwanted side effects in off-target tissues, such as immunosuppression and metabolic dysregulation, limit the agent’s therapeutic potential. Experimental methods to investigate this these questions are limited because genetic perturbation of mTOR does not recapitulate pharmacological inhibition. Therefore, here we describe selecTOR, a novel chemical-genetic strategy to target the activity of a Rapamycin analog to specific cell populations. We determined that the Rapamycin derivative Rapa*-3Z has greatly reduced ability to bind wild-type FKBP12 and inhibit mTOR in *S. cerevisiae*, mammalian cells, and *Drosophila*. We identified a single mutation in FKBP12 that restores Rapa*-3Z binding and rescues inhibition of mTOR. We showed that expression of mutant FKBP12 selectively sensitizes cells and tissues to inhibition by Rapa*-3Z. Finally, we used selecTOR to identify tissues that are responsible for the effects of TOR pathway inhibition on *Drosophila* larval development.

Previous bump-hole orthogonal Rapamycin analogs focused on reengineering the Rapamycin-FRB binding interface to prevent inhibition of native mTOR (34, 35). Adapting this system for tissue-specific targeting of mTOR would require tissue-specific knock-in or overexpression of mutant mTOR, potentially altering mTOR function in an unexpected manner. In contrast, our approach does not alter mTOR, which allows it to function as the wild-type kinase until the moment of inhibitor treatment. Additionally, mutant FKBP12 operates though a gain-of-function mechanism, so it can be simply over-expressed to sensitize the cells to inhibition, bypassing the need to install a mutation at the native FKBP12 locus.

By leveraging the specificity and modularity of genetics with the temporal control and therapeutic relevance of pharmacology, we hope to have built a tool that can reveal improved Rapamycin-based therapeutic strategies and create a clearer picture of mTOR’s fundamental role in regulating organismal biology through distinct cell populations. The application of selecTOR to animal models could address long unanswered questions about how mTOR contributes to disease and aging. For example, targeted inhibition of lymphocytes could determine whether Rapamycin’s immune-modulatory effects limit or contribute to its effects on aging. In addition, several mTOR pathway inhibitors have been shown to be effective at increasing lifespan and alleviating age-related disorders (36–38). However, a major drawback with these compounds is their lack of tissue-specificity. Our system circumvents this limitation and thus facilitates studies aimed at unravelling the complete therapeutic potential of this important pathway.

Once the relevant tissue targets have been identified, one could imagine applying a number of approaches to target specific tissues pharmacologically. Physicochemical properties can be tuned to modulate drug partitioning through natural barriers such as the blood-brain-barrier, as seen with second-generation antihistamines (39, 40). Drug conjugation can take advantage of endogenous receptors to achieve preferential uptake in specific tissues (41, 42). Additionally, the use of nanoparticles or other delivery methods can promote both passive and active uptake of a drug into a target tissue (43).

The chemical-genetic approach applied here to Rapamycin could be generalized to other “molecular glue” small molecules that rely on auxiliary proteins. The natural products FK506 and cyclosporin A depend on FKBP12 and Cyclophilin, respectively, to bind and inhibit Calcineurin (44), and PROTACs require binding to an E3 ligase to degrade their target protein (45). Making these molecules dependent on a mutant auxiliary protein would allow their activity to be targeted to specific cell populations, which may reveal new therapeutic modalities or uncover tissue-specific functions of their respective protein targets.

## Materials and Methods

### Chemical synthesis

All Rapa* compounds were prepared by similar procedure with respective hydroxylamine hydrochloride derivatives. Rapa*-3Z was prepared as follows. Rapamycin (50 mg, 0.055 mmol) and ethoxyamine hydrochloride (8 mg, 0.082 mmol) were dissolved in methanol (2 mL). Pyridine (6.6 μL, 0.082 mmol) was added by pipette and the reaction was stirred at room temperature for 24h. The crude material was purified by preparative reverse-phase HPLC (50-100% CH_3_CN in water containing 0.1% formic acid, RediSep C-18 prep column). *Z*-isomer product eluted before *E*-isomer. Desired fractions were combined and lyophilized to give the desired product as a white powder (36 mg, 69% yield). LC-MS (ESI+, [M+Na^+^]) *m/z* = 979.6. ^1^H NMR (400 MHz, Chloroform-*d*; Supplemental Figure S10).

### Yeast stocks and maintenance

BY4741 *fpr1::URA3* yeast were generated by homologous recombination, selected on Ura-synthetic complete media, and subsequently grown in YPD (46). Human FKBP12 coding sequence was cloned via Gibson assembly into a pCH043 yeast expression vector which had been previously modified with a HIS3 selection ORF. The C-terminal HA-tag and F46/V55 mutations were introduced by QuickChange PCR. Plasmids were transformed into the BY4741 *fpr1::URA3* strain and selected on His-synthetic complete media. Isolated strains were verified by plasmid sequencing. To prepare yeast protein lysates, 2 OD from an overnight culture was resuspended in 200 μl 0.1 M NaOH and incubated at room temperature for 15 minutes. Cells were pelleted, resuspended in 50 μL 2x SDS loading dye, and heated at 95°C for 5 minutes. Following centrifugation at 20,000 x g for 10 minutes, supernatant was saved for analysis.

### Disk diffusion assay

Yeast strains containing FKBP12-expression vectors were shaken overnight at 30°C in 3 mL His-synthetic complete media. 0.1 OD of culture was spun down, resuspended in 1 mL sterile water, and 100 μL was spread on His-synthetic media containing galactose/raffinose to induce protein expression. Plates were incubated at 30°C for 1h to dry. Filter paper circles were prepared with a hole punch, placed on the plate, and 5 μL of DMSO, 100 μM Rapamycin, or 100μM Rapa* compound was pipetted onto the filter paper. Plates were incubated at 30°C for 3 days before zones of growth inhibition were measured.

### Ternary complex formation

TR-FRET assay was performed using recommended conditions from Cisbio with slight modifications. FKBP12-HA, FKBP12 V55G-HA, and GST-mTOR_FRB_ recombinant proteins were each diluted in assay buffer (20 mM HEPES pH 7.5, 150 mM NaCl, 0.05% Tween-20, 0.1% BSA, 0.5 mM DTT) to 5 nM. Anti-GST-Tb (Cisbio) and Anti-HA-XL665 (Cisbio) were diluted to 0.625 μg/mL and 5 μg/mL, respectively, in assay buffer. Rapa* compounds and FK506 were diluted to 50 μM and Rapamycin to 5 μM in assay buffer containing 5% DMSO, and 3-fold serial dilution series were made from these solutions. 4 μL diluted FKBP12-HA protein, 4 μL diluted GST-mTOR_FRB_ and 4 μL of compound or DMSO control were mixed in a well of a black low-volume 384-well plate (Corning 4514) and the mixture was incubated at room temperature for 1h. 4 μL of Anti-GST-Tb and 4 μL Anti-HA-XL665 were then added, and the mixture was incubated at room temperature for an additional 1h. Time-resolved fluorescence was read on a TECAN Spark 20M plate reader with the following parameters: Lag time: 60 μs; Integration time: 500 μs; Read A: Excitation filter 320(25) nm, Emission filter 610(25) nm, Gain 130; Read B: Excitation filter 320(25) nm, Emission filter 665(8) nm, Gain 165. Three replicates were performed for each assay condition.

### X-ray crystallography

2 μL of 100 mM Rapa*-3Z in DMSO was added to 100μl of 1 mM FKBP12 V55G in 20 mM Tris pH 8.0 and incubated overnight at 4°C. Precipitate was removed by centrifugation at 20,000 x g for 10 minutes at 4°C. In a 15-well hanging drop plate with 500 μL well solution (0.1 M sodium tartrate, 18% PEG 3350), drops were set with 1 μL protein solution and 1 μL well solution and incubated at room temperature for 4 days. 10 μL cryo buffer (0.1 M sodium tartrate, 18% PEG 3350, 25% glycerol) was added to drops before crystals were looped and flash frozen in liquid nitrogen. Data were collected at Beamline 8.2.2 of the Advanced Light Source (LBNL, Berkeley, CA). Data were indexed and integrated using XDS (47). Molecular replacement was performed using Protein Data Bank 1FKB as a search model and the model was refined and built using PHENIX (48) and Coot (49). Structures were visualized with ChimeraX (50).

### Cell Culture

All cell lines were maintained in DMEM (Gibco) supplemented with 10% heat-inactivated FBS (Axenia Biologix) at 37°C in 5% CO_2_. Flp-In T-REx 293 cells were obtained from Thermo Fisher supplemented with 15 μg/mL blasticin and 100 μg/mL Zeocin. Flp-In T-REx 293 FKBP12 V55G cells were supplemented with 15 μg/mL blasticin and 150 μg/mL hygromycin B. FKBP12 V55G expression was induced with 1 μg/mL doxycycline 24 h prior to drug treatments. MCF7 cells were obtained from ATCC, and MCF7 mTOR F2108L cells were described previously (30). When indicated, cells were treated with drugs at 60-80% confluency at a final DMSO concentration of 0.1%. At the end of the treatment period, cells were placed on ice and washed once with PBS. Cells were scraped with a spatula and lysed in RIPA buffer supplemented with protease and phosphatase inhibitors (cOmplete and phosSTOP, Roche) on ice for 5 min. Lysates were clarified by high-speed centrifugation. Lysate concentration was determined by Bradford assay (Thermo Fisher) and adjusted to 2 mg/mL with additional RIPA buffer. Samples were mixed with 5x SDS loading dye and heated at 95°C for 5 min.

### Gel electrophoresis and Western blot

SDS-PAGE was run with Novex 4-12% Bis-Tris gel (Invitrogen) in MOPS running buffer at 200V for 40 minutes. Samples were loaded at 20 μg lysate per lane. Protein bands were transferred onto 0.2 μm nitrocellulose membranes (Bio-Rad) using wet tank transfer apparatus (Bio-Rad Criterion Blotter) in 1x TOWBIN buffer with 10% methanol at 75V for 45 min. Membranes were blocked in 5% BSA-TBST for 1h at room temperature. Primary antibody binding was performed with the indicated antibodies at manufacturer’s recommended dilution in 5% BSA-TBST at 4°C overnight. After three 5 min. washes with TBST, secondary antibodies (Li-COR) were added as solutions in 5% skim milk-TBST at 1:10,000 dilutions. Secondary antibodies were incubated for 1h at room temperature. Following three 5 min. washes with TBST, membranes were imaged on a Li-COR Odyssey fluorescence imager. Blot quantification was performed in Fiji and data were graphed and fit in Prism 9 (GraphPad). The following primary antibodies were obtained from Cell Signaling Technology (CST), Abcam (ab), and ProteinTech (PT): S6 (CST 2317), phospho-S6 S240/244 (CST 5364), S6K (CST 2708), phospho-S6K T389 (CST 9206), Akt (CST 2920), phospho-Akt S473 (CST 4060), HA (CST 3724), FKBP12 (ab58072), GAPDH (PT 60004).

### Cell viability

Cells were seeded into 96-well tissue-culture treated plates (Greiner) and were allowed to incubate for 24 hours. Cells were treated with indicated compounds and incubated for 72 h (100 μL final well volume, 0.1% DMSO). Cell viability was assessed by CellTiter-Glo (Promega). Assay reagent was diluted 1:4 in PBS containing 1% Triton X-100 and 100 μL was added to each well and mixed on a plate shaker for 20 minutes. Luminescence was measured on a Tecan Spark 20M.

### Fly stocks and maintenance

All *Drosophila* stocks and crosses were maintained on standard cornmeal-molasses food at 25°C using a 12-h light/dark cycle. To generate UAS-FKBP12 V55G-HA transgenic lines, human FKBP12 V55G-HA ORF was cloned into a pUAST-attB vector between EcoR-I and Xho-I restriction sites. This construct was then integrated into an attB site using phiC31 integrase with the assistance of Rainbow Transgenic Flies, Inc. The driver stocks *da-GAL4* (ubiquitous), *elav-GAL4* and *nSyb-GAL4* (neuronal-specific), *Mef2-GAL4* and *mhc-GAL4* (musculature), and *2-286-GAL4* and *phm-GAL4* (ring-gland-specific) were obtained from Bloomington Drosophila Stock Center. The crossing schemes used in this study are depicted in Supplemental Figure S11. For Rapamycin or Rapa*3Z related experiments, fly food was prepared freshly 24h before each experiment.

### Larval development experiments

For larval development assays, UAS-FKBP12 V55G-HA transgenic flies were set up for crosses with driver flies in freshly prepared fly food containing either Rapamycin or Rapa*-3Z (dissolved in ethanol). Fly food containing ethanol vehicle was used as control. All images were captured on day 9 after setting up the crosses unless otherwise mentioned, and pupae were counted on day 9. Three biological replicates were used for each experiment. Data were analyzed using Prism 9.0 (GraphPad).

## Acknowledgments

We thank Dr. Ziyang Zhang and Shizhong Dai for providing materials used in experiments, Dr. Qi Hu and Dr. Keelan Guiley for assistance with crystallography collection and analysis, and Dr. William Weiss, Dr. QiWen Fan, Dr. Nicole Nasholm, Dr. Daniel Schwarz, Taia Wu, and all members of the Shokat and Pallanck labs for their helpful input and discussion. This work was supported by the National Cancer Institute of the National Institutes of Health under awards 1R01CA221969 and 5F31CA243439, the National Institute on Aging of the National Institutes of Health under award R01AG057330, the Samuel Waxman Cancer Research Foundation, and the Howard Hughes Medical Institute. We thank reviewer #1 who suggested additional tissue-specific GAL4 driver lines to validate our initial results.

## Notes

### Competing Interest Statement

K.M.S. is an inventor on patents owned by UCSF covering mTOR targeting small molecules licensed to Calithera Biosciences and Revolution Medicines. K.M.S. has consulting agreements for the following companies, which involve monetary and/or stock compensation: Revolution Medicines, Black Diamond Therapeutics, BridGene Biosciences, Denali Therapeutics, Dice Molecules, eFFECTOR Therapeutics, Erasca, Genentech/Roche, Janssen Pharmaceuticals, Kumquat Biosciences, Kura Oncology, Mitokinin, Type6 Therapeutics, Venthera, Wellspring Biosciences (Araxes Pharma), Turning Point, Ikena, Initial Therapeutics and BioTheryX.

### Summary of Updates

Figure 6 and related text updated to include additional Drosophila driver lines. Conclusions regarding the role of the ring gland updated to reflect these new data. Supplemental files updated.

