## Supplemental Information for "Tissue-Restricted Inhibition of mTOR Using Chemical Genetics"

**This PDF file includes:**

Extended Methods

Supplemental Figures S1 to S11

Supplemental Table S1

**Extended Methods.**

*Recombinant protein expression and purification*

Human FKBP12 coding sequence was cloned a pET-47b bacterial expression vector using Gibson assembly and a C-terminal HA-tag was added by QuickChange PCR. FKBP12 proteins were expressed as 8xHis fusions in *E. coli* BL21 (DE3) cells. Cultures in Terrific Broth were grown to OD ~0.6, then protein expression was induced with 1 mM isopropyl b-D-1-thiogalactopyranoside (IPTG) at 18˚C overnight. Cells were lysed by microfluidizer in lysis buffer (20 mM Tris pH 8.0, 500 mM NaCl) containing protease inhibitor (cOmplete, Roche), centrifuged at 20,000 x g for 1 h, and purified by nickel chromatography including washes with lysis buffer containing 10 mM imidazole. Proteins were eluted using lysis buffer containing 250 mM imidazole. 2 units of HRV 3C protease (Takara Bio) were added and samples were dialyzed into storage buffer (20 mM Tris pH 8.0) overnight at 4˚C. Samples were then purified by size exclusion chromatography, concentrated to 10-20 mg/mL, and snap frozen in liquid nitrogen. pGEX-2T FRB was a gift from Jie Chen (Addgene plasmid #26607). mTOR FRB was expressed as a GST-fusion and purified as previously described (1).

*Generation of 293 Flp-In T-REx FKBP12 V55G cell line*

FKBP12 V55G-expressing Flp-In T-REx 293 cells were generated using recommended conditions from Thermo Fisher. Briefly, human FKBP12 V55G-HA coding sequence was cloned into the pcDNA5/FRT/TO targeting vector by Gibson assembly. Targeting vector was transfected into 293 Flp-In T-REx cells with the pOG44 vector at a 1:9 ratio using Lipofectamine LTX. 3 days after transfection, cells were split and selected with 150 µg/mL hygromycin B. Resulting colonies were combined into a single population.

*Generation of MCF7 FKBP12 V55G cell line*

Human FKBP12 V55G-HA coding sequence was cloned into a pWPI lentiviral vector at the PmeI site. Lentivirus was produced in HEK293T cells following transfection with standard packaging vectors using the TransIT-LT1 Transfection Reagent (Mirus Bio). Viral supernatant was collected 2 days after transfection, filtered through 0.44 µm PVDF syringe filter, and frozen before transduction. MCF7 cells were transduced with lentiviral supernatant and GFP-positive cells were isolated through FACS using a Sony SH800 cell sorter.

*PAMPA and Caco-2 membrane permeability assays*

PAMPA was performed by Quintara Discovery as described previously (2). Compounds were analyzed at 100 µM in PBS and measured using liquid chromatography-mass spectrometry (LC-MS) after 5 hours. Bidirectional Caco-2 cell permeability assay was performed by Quintara Discovery. Compounds were analyzed at 10 µM and measured by LC-MS after 1 hour.

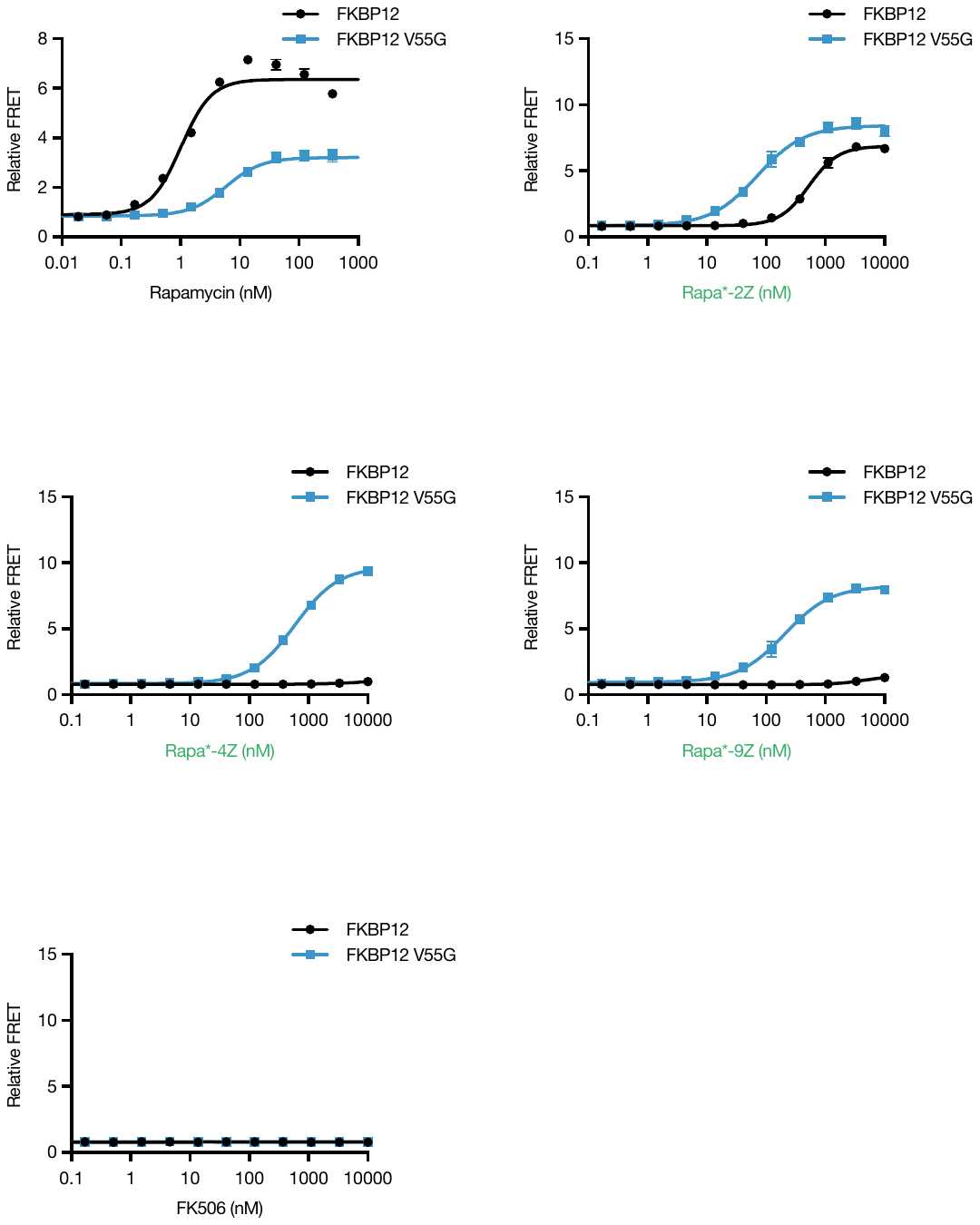

Supplemental Figure S1. Ternary complex formation with additional compounds. Ternary complex formation measured by TR-FRET. Rapamycin is included as a positive control, and FK506 is included as a negative control.

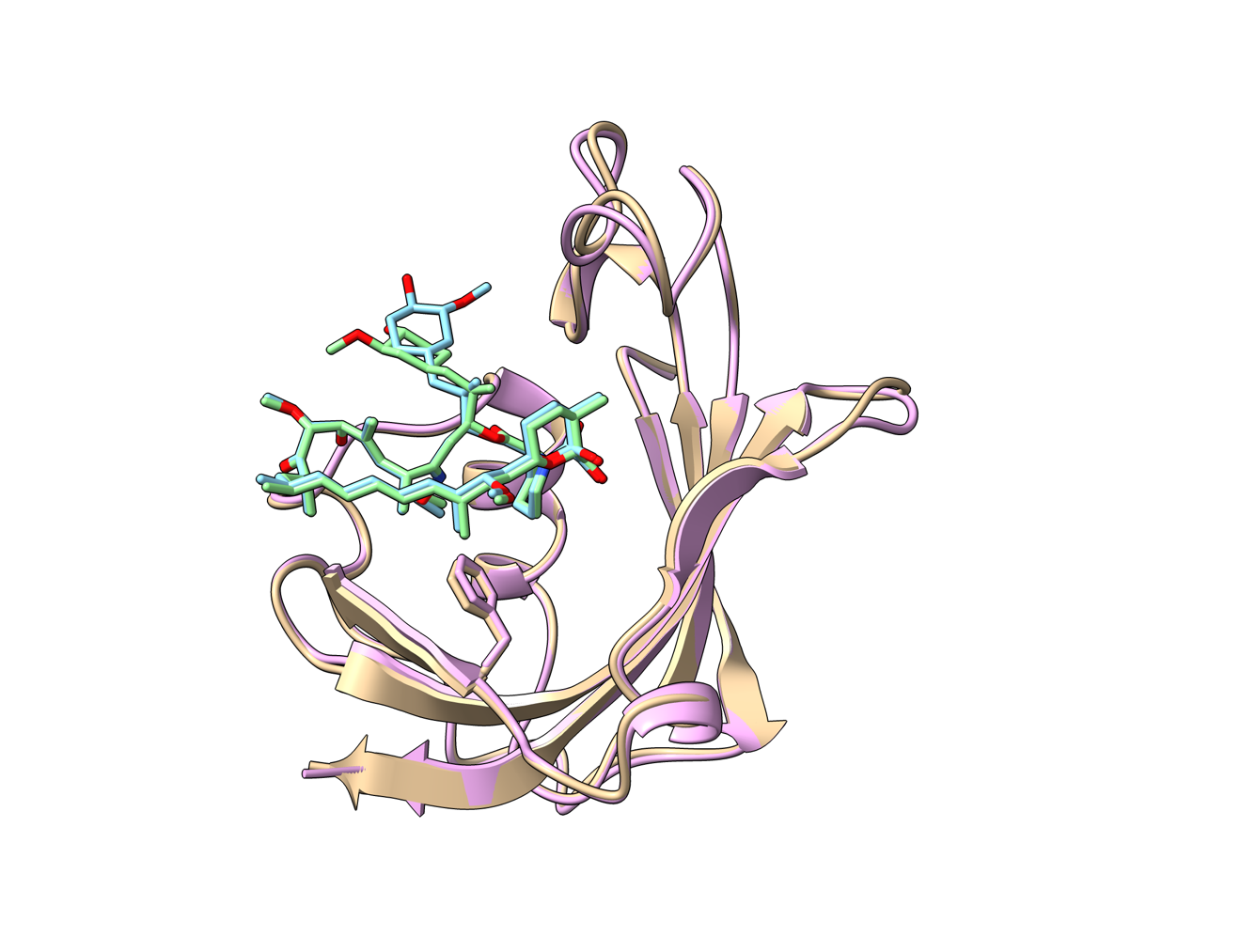

Supplemental Figure S2. Rapa*-3Z conformations in crystal structure. Chain A (tan, blue), chain B (purple, green).

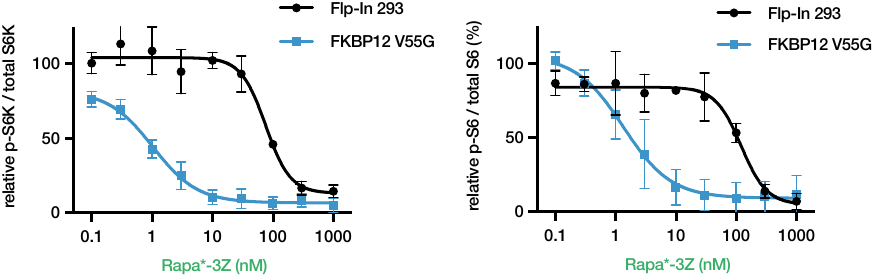

Supplemental Figure S3. Dose-dependent mTOR inhibition by Rapa*-3Z. Western blot quantification after 24 hour Rapa*-3Z treatment in Flp-In 293 cells. n = 2 for p-S6K and n = 3 for p-S6. Error bars show s.d.

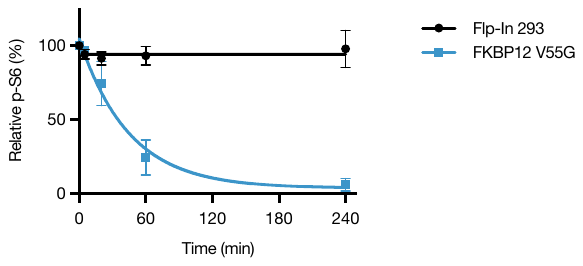

Supplemental Figure S4. Time course of mTOR inhibition by Rapa*-3Z. Quantification of p-S6 Western blot following Rapa*-3Z treatment in Flp-In 293 cells with or without over-expression of FKBP12 V55G. n = 3, error bars show s.d.

Supplemental Figure S5. Rapa*-3Z has no influence on larval development in control (w1118) flies. Vials showing pupae formed by w1118 control flies grown in the presence or absence of Rapa*-3Z (300 and 400 µM) 14 days after egg laying.

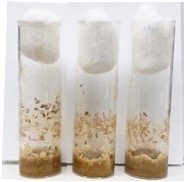

Untreated

Rapa*-3Z

(300 µM)

w1118 control flies

Rapa*-3Z

(400 µM)

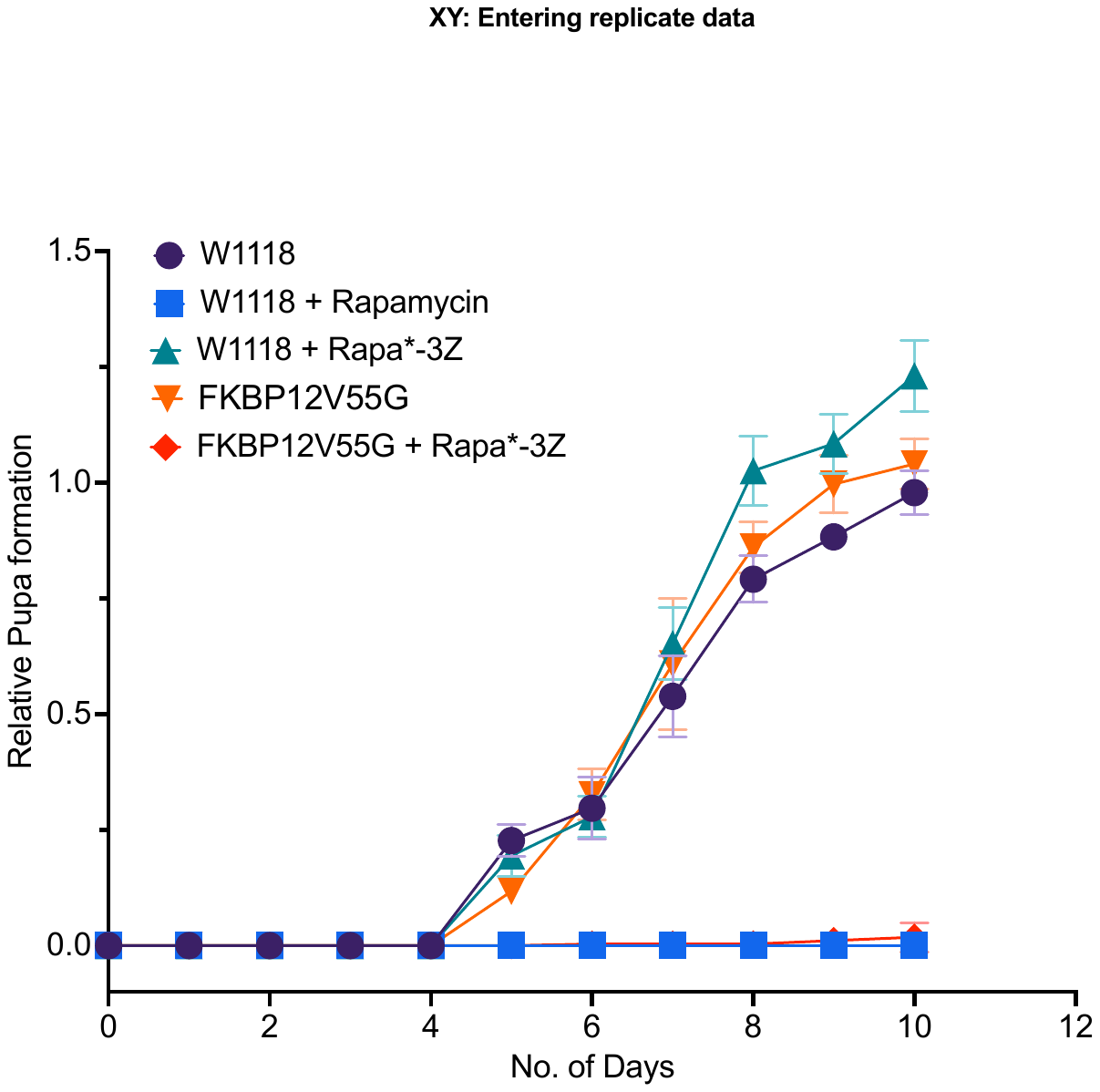
Supplemental Figure S6. Growth curve of larval development illustrating the developmental delay of transgenic animals. Plot shows relative number of pupae formed on the given number of days in flies of the given genotypes in the presence or absence of Rapamycin (100 µM) or Rapa*-3Z (100 µM). Error bars represent standard deviation from three biological replicates (n = 3).

|  | PAMPA | Caco-2 | |
| --- | --- | --- | --- |
|  | P_app_ | P_app_ (A -> B) | P_app_ (B -> A) |
| Rapamycin | 0.209 | 0.8 | 2.3 |
| Rapa*-1Z | 0.002 | < 0.3 | 1.6 |
| Rapa*-3Z | 0.053 | < 0.5 | 1 |
| Rapa*-9Z | 0.014 | < 0.1 | 0.4 |

**Supplemental Figure S7.** Membrane permeability of Rapamycin analogs. Apparent permeability (P_app_) reported in units of 10^-6^ cm/s.

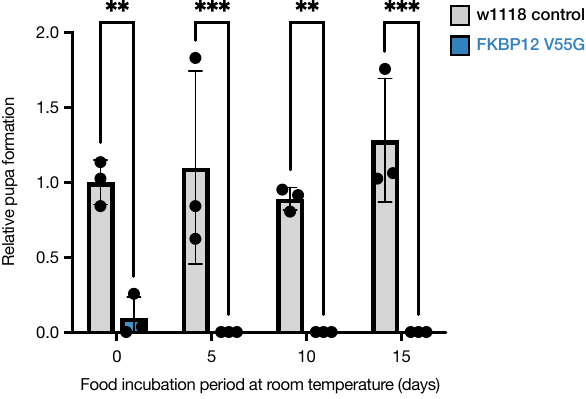

Supplemental Figure S8. Rapa*-3Z is stable in fly food for at least 15 days. Relative number of pupae was represented from indicated vials 9 days after egg laying in animals (w1118 control and FKBP12 V55G mutant) treated with 100 µM Rapa*-3Z. At least three independent experiments were performed (n=3). Significance was determined by two-way ANOVA multiple comparison test. * p < 0.05; ** p< 0.01; *** p < 0.001; **** p < 0.0001.

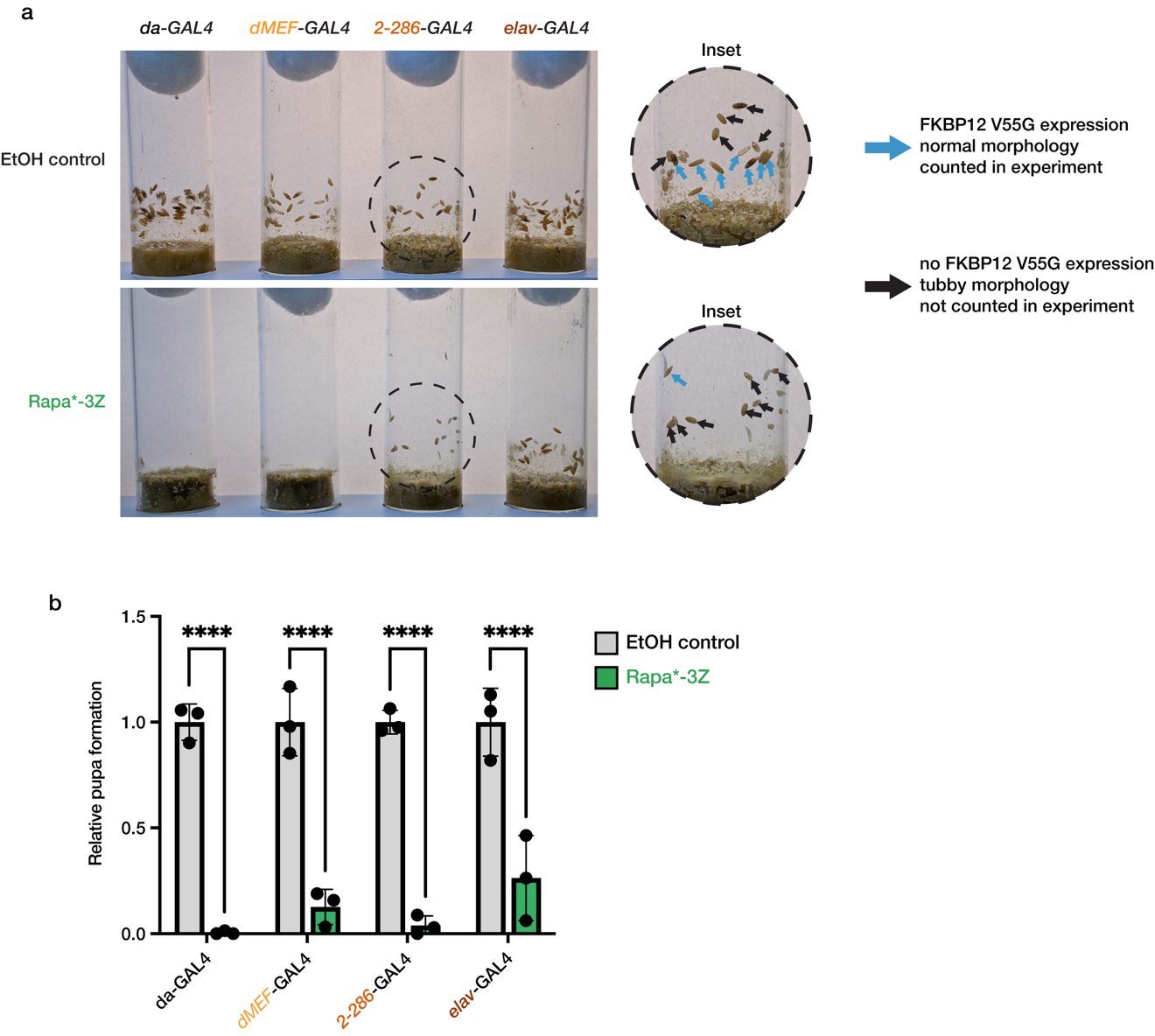

**Supplemental Figure S9.** Tissue-specific inhibition of TOR pathway showed larval developmental delays. (a) FKBP12 V55G was expressed using a ubiquitous GAL4 driver (*da-GAL4*), a muscle-specific driver (*dMEF-GAL4*), a ring-gland-specific driver (*2-286-GAL4*), and a nervous system-specific driver (*elav-GAL4*). Vials illustrate the number of animals of the indicated genotypes that develop to the pupal stage in the presence or absence of 100 µM Rapa*-3Z 9 days after egg laying. The *2-286-GAL4* insertion is lethal when homozygous, so flies bearing the 2-286 GAL4 driver in trans to the TM6 balancer were crossed to flies homozygous for the FKBP12 V55G transgene. The TM6 balancer chromosome contains the dominant Tb marker which confers a tubby phenotype (black arrow) that is characterized by a shorter and rounder pupal case. Pupae with the tubby phenotype do not drive FKBP12 V55G in the ring gland and were therefore not counted in the experiment. Pupae with normal morphology (blue arrow) do drive FKBP12 V55G expression in the ring gland and were counted in the experiment. (b) The relative number of pupae detected 9 days after egg laying in animals of the indicated genotypes treated with 100 µM Rapa*-3Z (n=3). Tubby pupae not included in analysis of the *2-286-GAL4* driver. Significance was determined by group comparison via two-way ANOVA *p < 0.05; ** p < 0.001; *** p = 0.0001; **** p < 0.0001

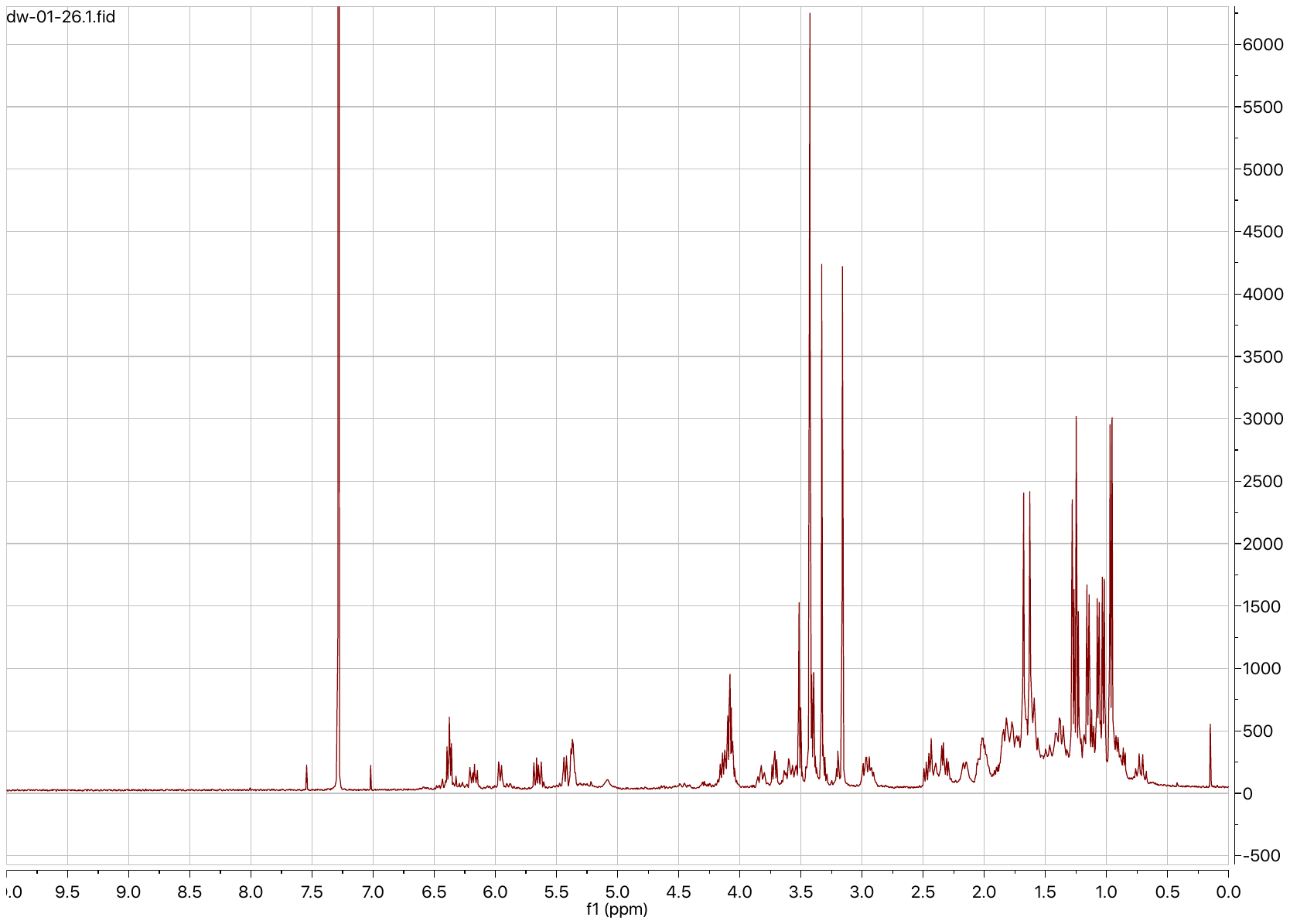
 **Supplemental Figure S10.** Rapa*-3Z ^1^H-NMR spectrum in CDCl_3_.

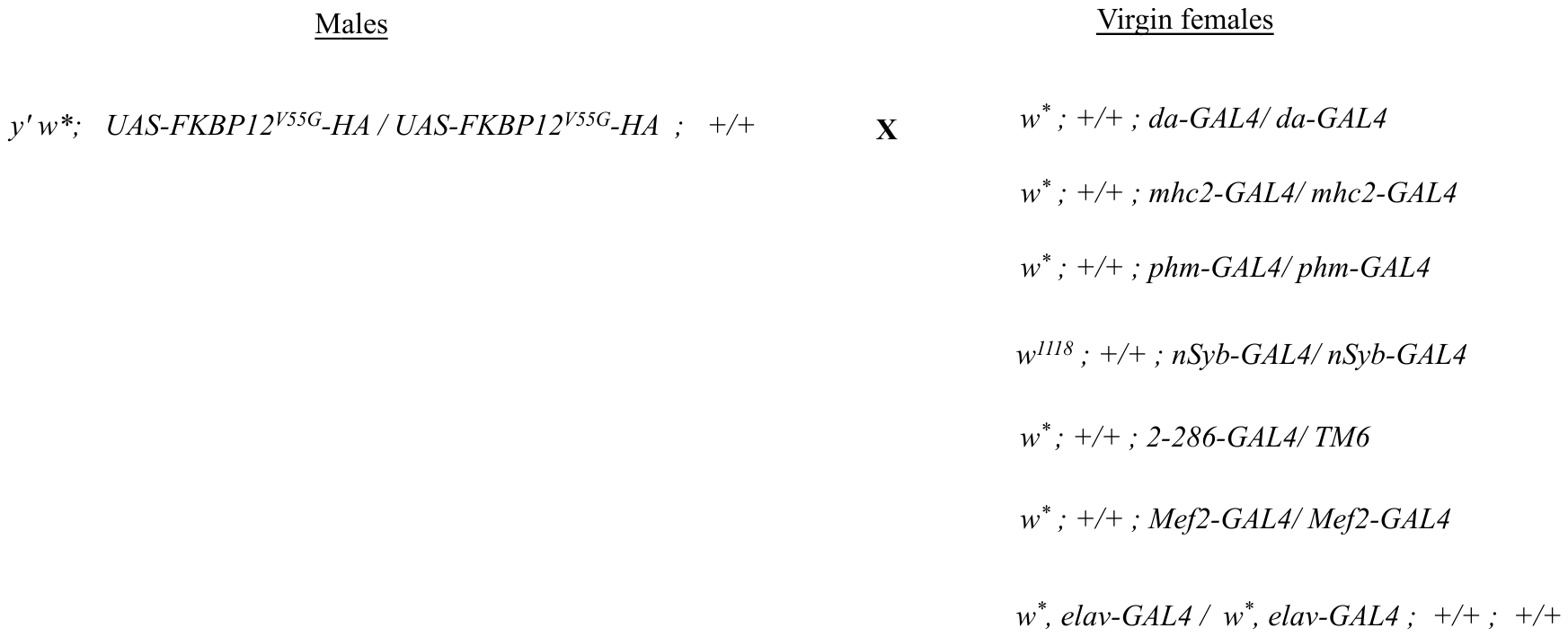

**Supplemental Figure S11.** Crossing schemes used to generate mutant flies expressing FKBP12 V55G.

Supplemental Table S1. Crystallographic data collection and refinement statistics

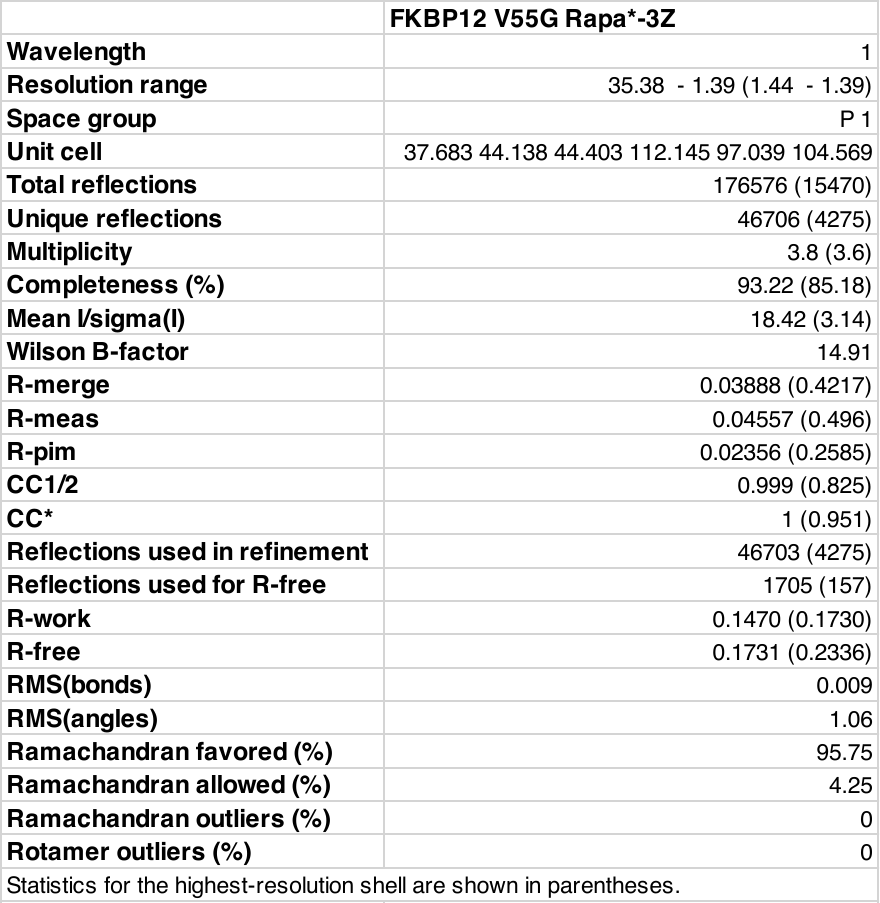
